# Nonlinear dynamics in auditory cortical activity reveal the neural basis of perceptual warping in speech categorization

**DOI:** 10.1101/2021.12.07.470603

**Authors:** Jared A. Carter, Eugene H. Buder, Gavin M. Bidelman

## Abstract

Surrounding context influences speech listening, resulting in dynamic shifts to category percepts. To examine its neural basis, event-related potentials (ERPs) were recorded during vowel identification with continua presented in random, forward, and backward orders to induce perceptual nonlinearities. Behaviorally, sequential order shifted listeners’ categorical boundary vs. random delivery revealing perceptual warping (biasing) of the heard phonetic category dependent on recent stimulus history. ERPs revealed later (∼300 ms) activity localized to superior temporal and middle/inferior frontal gyri that predicted listeners’ hysteresis magnitudes. Findings demonstrate that top-down, stimulus history effects on speech categorization are governed by interactions between frontotemporal brain regions.

## 1. INTRODUCTION

In speech perception, listeners group similar sensory cues to form discrete phonetic labels—the process of categorical perception (CP). Spectral features vary continuously, but reducing acoustic cues to discrete categories enables more efficient use of speech sounds for linguistic processing^1,2^. The extent to which phonetic speech categories built from acoustic-sensory cues are influenced by perceptual biasing (top-down influences) has been debated. On one hand, categories might arise due to innate psychophysiological constraints^3^. Alternatively, there is ample evidence that top-down processing influences speech categorization as suggested by enhancements observed in highly proficient listeners^4-7^ and biasing effects, when individuals hear a different category depending on the surrounding speech context^8^.

Changes in auditory-perceptual categories due to stimulus history are a form of nonlinear dynamics. Nonlinear dynamics in CP are especially prominent at the perceptual boundary, where different patterns of behavioral identification can result for otherwise identical speech sounds: hysteresis (i.e., percept continuing in the same category beyond the theoretical boundary) or enhanced contrast (i.e., percept changing to the other category before the theoretical boundary)^9-11^. Both stop consonant and vowel continua can produce context-dependent shifts in perception, though stronger perceptual warping occurs with more ambiguous speech sounds^12^.

Event-related brain potentials (ERPs) have been used to examine the neural underpinnings of speech categorization^13-15^. ERPs reveal the brain performs its acoustic-to-phonetic conversion within ∼150 ms and differentiates even the same speech sounds when categorized with different perceptual labels^13^. Yet it remains unknown how neural representations of categories change with recent state history as seen in hysteresis and other perceptual nonlinearities inherent to speech perception^10^. Shifting percepts near a categorical boundary due to presentation order (i.e., how stimuli are sequenced) should yield measurable neural signatures if speech perception is indeed warped dynamically.

Here, we evaluated the effects of nonlinear dynamics on speech categorization and its brain basis. We aimed to resolve whether perceptual hysteresis in CP occurs at early (i.e., auditory-sensory) or later (i.e., higher-order, linguistic) stages of speech analysis. We measured behavioral and multichannel EEG responses during rapid phoneme identification tasks where tokens along an identical continuum were presented in random vs. serial (forward or backward) order. Based on previous studies examining nonlinear dynamics^9,10^ and top-down influences in speech CP^4,5,7^, we hypothesized (1) the location of listeners’ perceptual boundary would shift according to the direction of stimulus presentation and (2) perceptual warping would be accompanied by late modulations in the ERPs.

## 2. MATERIALS & METHODS

### 2.1 Participants

The sample included N=15 young participants (23.3 ± 3.9 years; 5 females) averaging 16.7±3.4 years of education. All spoke American English, had normal hearing (≤20 dB HL; 250– 8000 Hz), minimal musical training (≤3 years; average = 1.0 ± 1.3 years), and were mostly right-handed (mean = 75% ± 40% laterality)^16^. Each gave written informed consent in compliance with the University of Memphis IRB.

### 2.2 Stimuli & task

We used a 7-step vowel continuum from /u/ to /α/. Each 100 ms token had a fundamental frequency of 100 Hz (i.e., male voice). Adjacent tokens were separated by equidistant steps in first formant (F1) frequency spanning from 430 (/u/) to 730 Hz (/α/). We selected vowels over consonant-vowel (CV) syllables because pilot data suggested vowels were more prone to nonlinear perceptual effects (see **Figure S1**). We delivered stimuli binaurally through insert earphones (ER-3A) at 76 dB_A_ SPL. Sounds were controlled by MATLAB coupled to a TDT RP2 signal processor (Tucker-Davis Technologies, Alachua, FL).

There were three conditions based on how tokens were sequenced: (1) random presentation, and two sequential orderings presented serially between continuum endpoints and F1 frequencies (2) forward /u/ to /α/, 430 Hz to 730 Hz, and (3) backward /α/ to /u/, 730 to 430 Hz). Forward and backward directions on such a continuum were expected to produce perceptual warpings (i.e., hysteresis)^10^. Random and serial order conditions were presented in different blocks (randomized between participants). We allowed breaks between blocks to avoid fatigue.

Within each condition, listeners heard 100 presentations of each vowel (total = 700 per block). On each trial, listeners rapidly reported which phoneme they heard with a binary keyboard response (“u” or “a”). Following their response, the interstimulus interval was jittered randomly between 800 and 1000 ms (20 ms steps, uniform distribution).

### 2.3 Behavioral data analysis

#### 2.3.1 Psychometric function analysis

Identification scores were fit with sigmoid P = 1/[1 + e^−β1(x−β0)^], where P is the proportion of trials identified as a given vowel, x is the step number along the continuum, and β0 and β1 are the location and slope of the logistic fit estimated using non-linear least-squares regression^14,17^. Leftward/rightward shifts in *β0* location for the sequential vs. random stimulus orderings would reveal changes in the perceptual boundary characteristic of perceptual nonlinearity^10^. These metrics were analyzed using a one-way mixed-model ANOVA (subjects = random factor) with a fixed effect of condition (3 levels: random F1, forward F1, backward F1) and Tukey-Kramer adjustments for multiple comparisons. Reaction times (RTs) were computed as the median response latency for each token per condition. RTs outside of 250-2000 ms were considered outliers (i.e., guesses or attentional lapses) and were excluded from analysis^13,14^. RTs were analyzed using a two-way, mixed model ANOVA (subjects = random) with fixed effects of condition (3 levels: random F1, forward F1, backward F1) and token (7 levels).

#### 2.3.2 Cross-classification analysis of behavioral response sequences

To determine the effect of sequential presentation order (i.e., forward vs. backward F1) on behavioral responses, we performed cross-classification analysis on single-runs of the identification data (i.e., responses from tokens 1 – 7 or 7 – 1) in the Generalized Sequential Querier program (v 5.1.23; Mangold International https://www.mangold-international.com/en/products/software/gseq). This compared listeners’ category labels to specific tokens (e.g., trials where token 3 was labeled as “u” vs. “a”) when the stimulus continuum was presented in the forward (i.e., rising F1) versus backward (i.e., falling F1) direction. Biasing due to presentation order was quantified using Yule’s Q, an index of standardized effect size transformed from an odds ratio to vary from -1 to 1 which is superior to the odds ratio in being relatively unskewed, affording more direct statistical analysis^18^. In the current application, a Q of -1 means “u” selected more in the forward F1 condition and “a” selected more in the backward F1 condition; a Q of +1 indicates the opposite pattern; and values effectively equal to 0 indicate presentation order had no effect on response selection. This analysis allowed us to determine whether the direction of stimulus presentation (i.e., increasing/decreasing F1) shifted listeners’ category labels towards one endpoint of the continuum or the other (i.e., evidence of perceptual hysteresis). The non-0 responses at Tk3/Tk5 were used to classify participants as “hysteresis” vs. “enhanced contrast” listeners (i.e., those showing late vs. early biasing in their category labeling). See Table S1 for details.

### 2.4 EEG recording procedures and analysis

#### 2.4.1 EEG recording

Continuous EEGs were recorded during the speech identification task from 64 sintered Ag/AgCl electrodes at standard 10-10 scalp locations (NeuroScan Quik-Cap array)^19^. Continuous data were sampled at 500 Hz (SynAmps RT amplifiers; Compumedics NeuroScan) with an online passband of DC-200 Hz. Electrodes placed on the outer canthi of the eyes and superior/inferior orbit monitored ocular movements. Contact impedances were <10 kΩ. During acquisition, electrodes were referenced to an additional sensor placed ∼1 cm posterior to the Cz channel. Data were common average referenced for analysis.

#### 2.4.2 Cluster-based permutation analysis

To reduce data dimensionality, ”super-channel” clusters were computed by averaging adjacent electrodes over 5 left/right frontocentral scalp areas (see **Fig. 3**)^14,20^. We used cluster-based permutation statistics^21^ implemented in BESA® Statistics 2.1 to determine whether super-channel ERP amplitudes differed with presentation order. This ran an initial *F*-test across the whole waveform (i.e., -200–800 ms), contrasting random, forward, and backward F1 conditions. This step identified time samples and super-channels where neural activity differed between conditions (p < 0.05). Critically, BESA corrects for multiple comparisons across space and time. This was then followed by a second level analysis using permutation testing (N=1000 resamples) to identify significant post hoc differences between pairwise stimulus conditions (i.e., random/forward/backward stimulus orderings). Contrasts were corrected with Scheffe’s test using Bonferroni-Holm adjustments. Lastly, we repeated this analysis for tokens 3-5, representing stimuli surrounding the categorical boundary where warping was most expected.

#### 2.4.3 Distributed Source Analysis

We used Classical LORETA Analysis Recursively Applied (CLARA) distributed imaging with a 4 shell ellipsoidal head model (conductivities of 0.33 [brain], 0.33 [scalp], 0.0042 [bone], and 1.00 [cerebrospinal fluid]) on the difference wave to determine the intracerebral sources that account for perceptual non-linearities in speech categorization^22^. Source images were computed at a latency of 320 ms, where the scalp ERPs differentiated stimulus order based on the cluster-based statistics (see **Fig. 4**). Correlations between changes in β0 and CLARA activations evaluated which source regions predicted listeners’ perceptual warping of speech categories.

## 3. RESULTS

### 3.1 Behavioral data

#### 3.1.2 Psychometric function data

Listeners perceived vowels categorically in all three presentation orderings as evidenced by their sigmoidal identification functions (**Fig. 1A**). Slopes varied with presentation order (*F*_*2,28*_ = 6.96, *p* = 0.0463); this was driven by the forward condition producing stronger categorization than random (p = 0.0364) (**Fig. 1C**). The categorical boundary did not appear to change with condition when analyzed at the group level (F_2,28_ = 1.78, p = 0.1875) (**Fig. 1D**).

**Figure 1:**
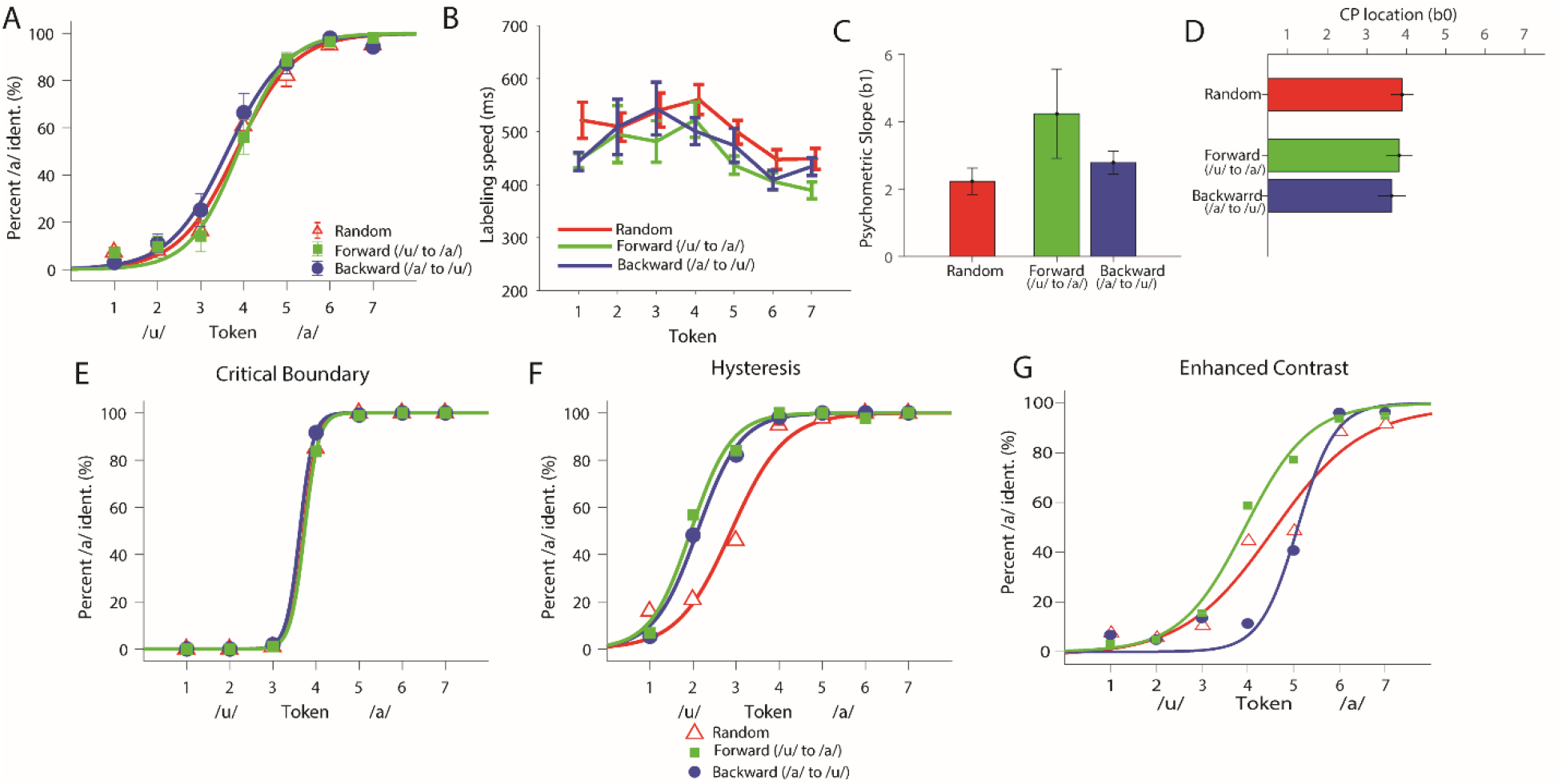
Behavioral speech categorization is modulated by stimulus presentation order revealing nonlinearities in perception. **(A)** Perceptual psychometric functions for phoneme identification when continuum tokens are presented in random vs. serial (forward: /u/→/α/ vs. backward: /α/→/u/) order. **(B)** Reaction times for speech identification. Sequential presentation (i.e., forward and backward) led to faster speech labeling speeds than random presentation. **(C)** Psychometric function slope was steeper for forward compared to random presentation. **(D)** Boundary location did not vary at the group level (cf. individual differences; Fig. 2). Individual differences reveal unique forms of perceptual nonlinearity across sub-classes of listeners (n=3 representative subjects). **(E)** Critical boundary listener, where the individual selects the same response regardless of presentation order. **(F)** Hysteresis listener, where the prior percept continues beyond the expected perceptual boundary (midpoint) as measured in sequential presentation (cf. panel E). **(G)** Enhanced contrast listener, where the category response flips earlier than expected during sequential presentation. See also Table 1 for classifications of these listeners. Error bars = ±1 s.e.m.

RTs varied with presentation order (F_2,292_ = 8.45, p = 0.0003) and token (F_6,292_ = 10.85, p < 0.0001) (**Fig. 1B**). Participants’ labeling was slower for random compared to forward (p = 0.0002) and backward (p = 0.0419) presentation orders. RTs were also slower nearly the continuum midpoint vs. endpoints (Tk4 vs. Tk1/7: p < 0.0001), consistent with previous studies demonstrating slower RTs for category-ambiguous speech sounds^7,14^,23,24. Pooling orders, comparisons between the left/right sides of the continuum (Tk1,2,3 vs. Tk5,6,7) indicated listeners responded to “α” vowels faster than “u” vowels (p < 0.0001). This suggests sequential presentation of the continua, regardless of direction, improved speech categorization speeds.

Despite limited changes in boundary location at the group level (**Fig. 1D**), perceptual nonlinearities were subject to stark individual differences (**Fig. 1E-G**). Some listeners were consistent in their percept of individual tokens regardless of presentation order (i.e., “critical boundary” response pattern); others persisted with responses well beyond the putative category boundary at continuum midpoint (i.e., hysteresis); and other listeners changed responses earlier than expected (i.e., enhanced contrast). Response patterns were, however, highly stable *within* listener; a split-half analysis showed *β0* locations were strongly correlated between the first and last half of task trials (*r=*0.86, *p* < 0.0001). This suggests that while perceptual nonlinearities (i.e., *β0* shifts) varied across listeners, response patterns were highly repeatable within individuals.

We performed further cross-classification analysis to characterize these individual differences in categorization nonlinearities. **Table S1** shows participants’ Yule’s Q values Tk3/5 (i.e., tokens flanking the β0), and, thus, their predominant “mode” of hearing the speech continua. Individuals with negative Qs showed hysteresis response patterns (N = 9), while those with positive Qs showed enhanced contrast patterns in perception (N = 4). Still others (N = 2) did not show perceptual nonlinearities and demonstrated neither hysteresis nor enhanced contrast.

### 3.2 Electrophysiological data

Figure 2 shows scalp ERP “super channels” to token 4 (critical stimulus at the perceptual boundary) across presentation orders (see Fig. S2 for raw ERP data). Cluster based permutation tests^21^ also revealed nonlinear (stimulus order) effects emerging ∼320 ms after speech onset, localized to left temporal areas of the scalp (omnibus ANOVA; p=0.03; **Fig. 3A**, shading). Post hoc contrasts revealed order effects were driven by larger neural responses for the random vs. forward F1 condition (p = 0.003). CLARA source reconstruction localized this nonlinear effect (i.e., ERP_random@Tk4_ > ERP_forward@Tk4_) to underlying brain regions in bilateral superior temporal gyri (STG) and medial (MFG) and inferior (IFG) frontal gyri (**Fig. 3B)**.

**Figure 2:**
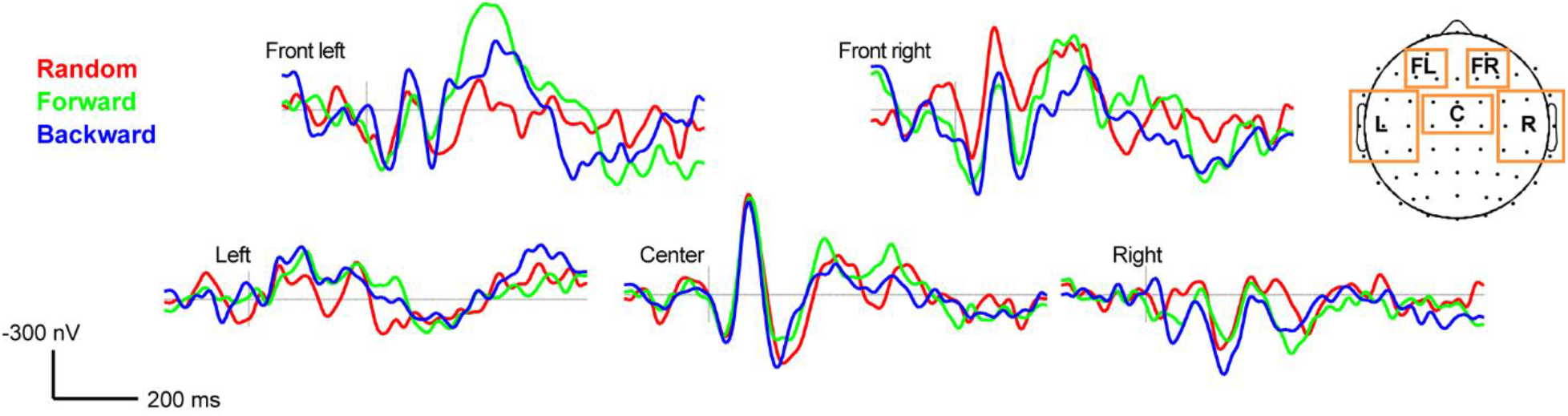
Grand average ERPs (at Tk4 = critical boundary stimulus) for forward, backward, and random presentation order of the vowel continuum. Boxes = “super-channel” electrode clusters. Negative voltage plotted up.

**Figure 3:**
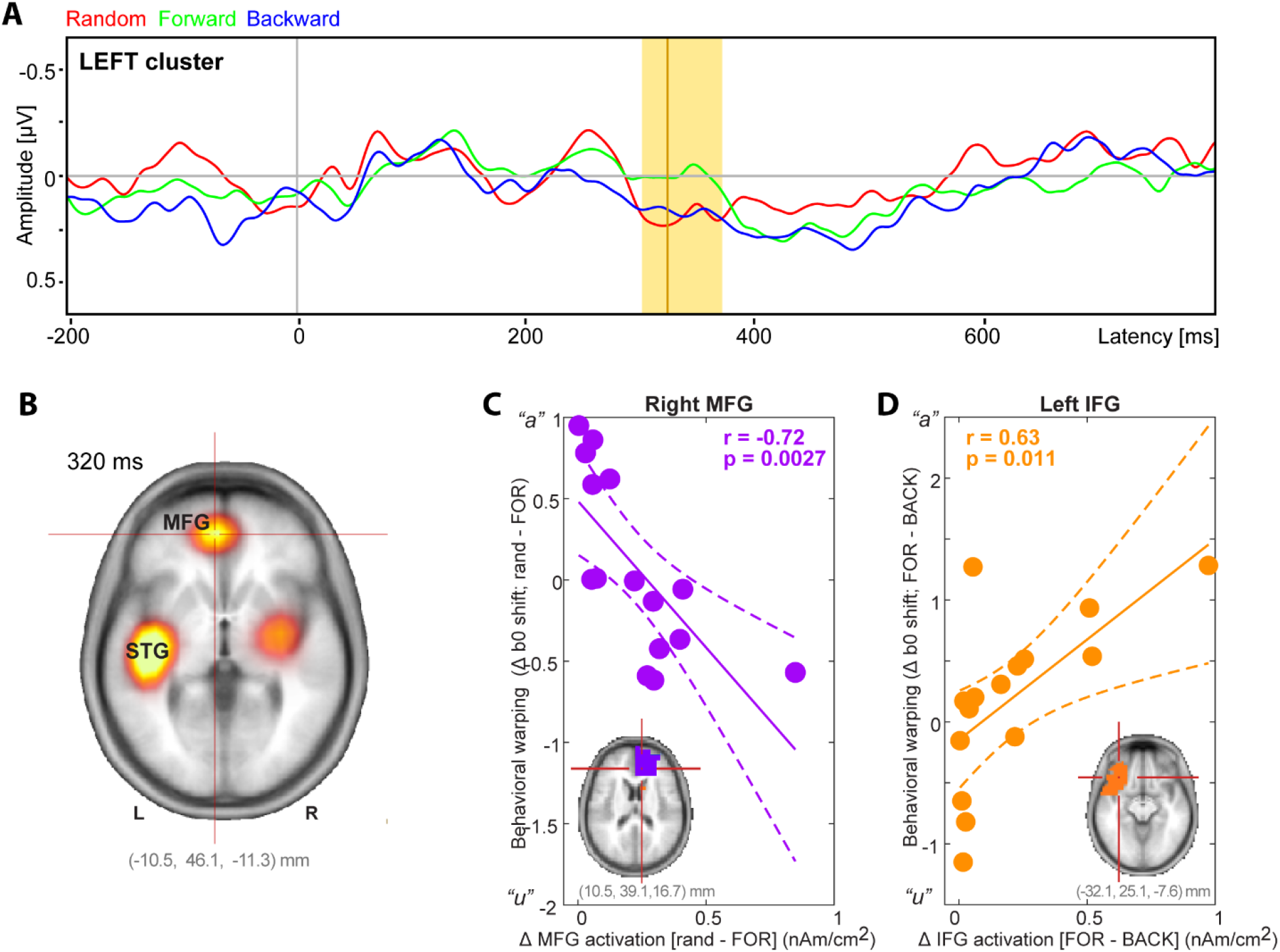
Perceptual nonlinearities in the auditory cortical ERPs emerge by ∼320 ms via interplay between frontotemporal cortices. (**A**) Cluster based permutation statistics contrasting responses to the identical Tk4 (stimulus at the continuum’s midpoint) in random, backward, and forward conditions. Nonlinearities in speech coding emerge by ∼300 ms (highlighted region) in the left super-channel. Line =maximal difference (322 ms). Negative = up. **(B)** CLARA source imaging contrasting the difference in activations to Tk4 during random vs. forward conditions. Nonlinearities in perceptual processing localize to bilateral superior temporal gyri and medial/inferior frontal gyri. (**C-D**) Brain-behavior correlations between the change in regional source activations and magnitude of hysteresis effect. Changes in right rMFG contrasting “randomness” (i.e., Δrandom - forward) are negatively associated with shifts in the CP boundary. Contrastively, modulations in left IFG contrast the *direction* of serial ordering (i.e., Δforward – backward) and are positively related to behavior.

We assessed the behavioral relevance of these neural findings via correlations between regional source activations (i.e., CLARA amplitudes @ 320 ms; **Fig. 3C-D**) and listeners’ behavioral CP boundary locations (*β0*). We found modulations in right MFG and left IFG with stimulus order were associated with behavioral *β0* shifts characteristic of perceptual warping but in opposite directions. Listeners with increased rMFG activation from random vs. ordered (forward) stimulus presentation showed lesser movement of their perceptual boundary [Pearson’s *r=*-0.72, *p*=0.0027]. In contrast, those with increased left IFG activation contrasting stimulus direction (i.e., Δ forward vs. backward) showed larger movement in *β0* location [*r=*0.63, *p*=0.011]. STG activations did not correlate with behavior (corrected *p*’s > 0.05).

## 4. DISCUSSION

By measuring EEG to acoustic-phonetic continua presented in different contexts (random, serial orderings), our data expose the brain mechanisms by which listeners assign otherwise identical speech tokens to categories depending on context. Behaviorally, perceptual nonlinearities were more prominent for vowels compared to CVs and were subject to stark individual differences. Behavioral warping corresponded with neural effects emerging ∼300 ms over left hemisphere with underlying sources in a frontotemporal circuit (bilateral STG, right MFG, left IFG). Our findings reveal stimulus presentation order strongly influences the neural encoding of phonemes and suggest that sequential warpings in speech perception emerge from top-down, dynamic modulation of early auditory cortical activity via frontal brain regions.

### Perceptual nonlinearities in categorization are stronger for vowels than CVs

We found vowels elicited stronger perceptual warping (i.e., changes in the CP boundary) than CV tokens. Vowels are generally perceived less categorically than CVs^12,25-27^. With the vowel state space already being more flexible than consonants, listeners are more free to alter perception based on prior history of other vowels. Formant frequencies intrinsic to vowels are relatively continuous in their variations, but also static. In contrast, formant transitions in CVs allow frequency comparisons within the stimulus itself^28,29^. Vowel percepts are thus more ambiguous categorically, and consequently more susceptible to contextual influences and individual differences^30^. Indeed, we find the magnitude and direction of perceptual warping strongly varies across listeners, consistent with prior work on perceptual hysteresis in both the auditory and visual domains^10,31^.

### Perceptual warping of categories is subject to stark individual differences

Behaviorally, we found minimal group-level differences in psychometric functions, with only an increase in slope when in the forward /u/ to /α/ direction versus random presentation. A change in identification slope indicates sequential presentation led to more abrupt category changes. The reason behind this direction-dependent effect is unclear but could be related to differences in perceptual salience between continuum endpoints. We can rule out differences due to vowel loudness as both /u/ and /α/ endpoints had nearly identical loudness according to ANSI S3.4 (2007) (/α/ = 71.9 phon; /u/ = 71.2 phon)^32^. Alternatively, /α/ might have been heard as being a more prototypical vowel (i.e., perceptual magnet)^33^. Conversely, RTs were faster in sequential compared to random presentation orders. RTs demonstrate the speed of processing, which increases (i.e., slows down) for more ambiguous or degraded tokens^7,29^ and decreases (i.e., speeds up) for more prototypical tokens^23^. Faster RTs during sequential presentation suggest a quasi-priming effect whereby responses to adjacent tokens were facilitated by the preceding (phonetically similar) stimulus.

Behavioral changes in category boundary location were most evident at the individual rather than group level (cf.^8,34^) and when speech tokens were presented sequentially. These findings suggest stimulus history plays a critical role in the current percept of phonemes. Listeners demonstrated three distinct response patterns (Table S1; hysteresis, enhanced contrast, critical boundary), differences which were largely obscured at the group level. This is consistent with previous work demonstrating trial-by-trial differences in nonlinear dynamics of speech categorization^9-11^. Critically, response patterns were highly stable *within* individuals, suggesting listeners have a dominant response pattern and/or apply different decision strategies (cf. biases) during categorization. This latter interpretation is also supported by the different regional activation patterns and their behavioral correlations. It is also reminiscent of lax vs. strict observer models in signal detection frameworks where, for suprathreshold stimuli, listeners’ response selection is primarily determined by their internal bias (i.e., preference for tokens at one end of the continuum)^35^.

### Electrophysiological correlates of perceptual warping

ERPs revealed late (∼320 ms post-stimulus) differences in response to token 4 (i.e., categorical boundary) between forward and random conditions over the left hemisphere. Sound-evoked responses in auditory cortex typically subside after ∼250 ms^36,37^. This suggests the stimulus order effects observed in our speech ERPs likely occur in higher-order brain regions subserving linguistic and/or attentional processing. The leftward lateralization of responses also suggests context-dependent coding might be mediated by canonical language-processing regions (e.g., Broca’s area)^38^. Indeed, source analysis confirmed engagement of extra-auditory brain areas including IFG and MFG whose activations scaled with listeners’ perceptual shifts in category boundary. In contrast, auditory STG, though differentially active during perceptual warping, did not correlate with behavior, *per se*.

Beyond its established role in speech-language processing, left IFG is heavily involved in category decisions, particularly under states of stimulus uncertainty (i.e., randomness, noise)^7,30^,39. Related, we find *direction-related* modulations in the perceptual warping of speech categories (to an otherwise identical sound) are predicted by left IFG engagement. This is consistent with notions that frontal brain regions help shape behavioral category-level predictions at the individual level^40^. Contrastively, rMFG correlated with changes in behavior between random vs. forward stimulus presentation, a contrast of ordered vs. unordered sequencing. MFG regulates behavioral reorienting and serves to break (i.e., gate) attention during sensory processing^41^. Additionally, it is active when holding information in working memory, such as performing mental calculations^42^, and has been implicated in processing ordered numerical sequences and counting^43^. The observed perceptual nonlinearities induced by serial presentation might therefore be driven by such buffer and comparator functions of rMFG as listeners hold prior speech sounds in memory and compare present to previous sensory-memory traces. In contrast, un-ordered speech presented back-to-back would not load such operations and thus may explain the reduced rMFG activity for random presentation. The simultaneous activation of canonical auditory areas (STG) concurrent with these two frontal regions leads us to infer that while auditory cortex is sensitive to category structure (present study; ^7,39^), top-down modulations from frontal lobes dynamically shapes category percepts online during speech perception.

## Acknowledgements

Work supported by the National Institute on Deafness and Other Communication Disorders (R01DC016267). Requests for data and materials should be directed to G.M.B.

