## Supplemental Material for "Nonlinear dynamics in auditory cortical activity reveal the neural basis of perceptual warping in speech categorization"

**Pilot experiment: Characterizing perceptual warping for vowels vs. CVs**

We first examined whether perceptual warping of categories varies among different speech sounds (vowels vs. consonants). To this end, we ran a pilot sample that included N=5 young adults. Participants were native speakers of American English and reported normal hearing. We used 7-step continua of vowels (/u/ to /ɑ/) with F1 frequencies spanning from 430 to 730 Hz and consonant-vowel (CV) syllables (/dɑ/ to /gɑ/) used in previous studies[7,44,45]. Listeners were instructed to listen to these stimuli through headphones and respond by clicking on an onscreen button whether they heard “oo” or “ah” in the vowel conditions and “da” or “ga” in the CV condition. With each condition, listeners heard 10 repetitions of each token (total = 70 tokens per condition). The pilot task was conducted via internet-based data collection using paradigms coded in E-Prime 3.0 delivered using E-Prime Go[46].

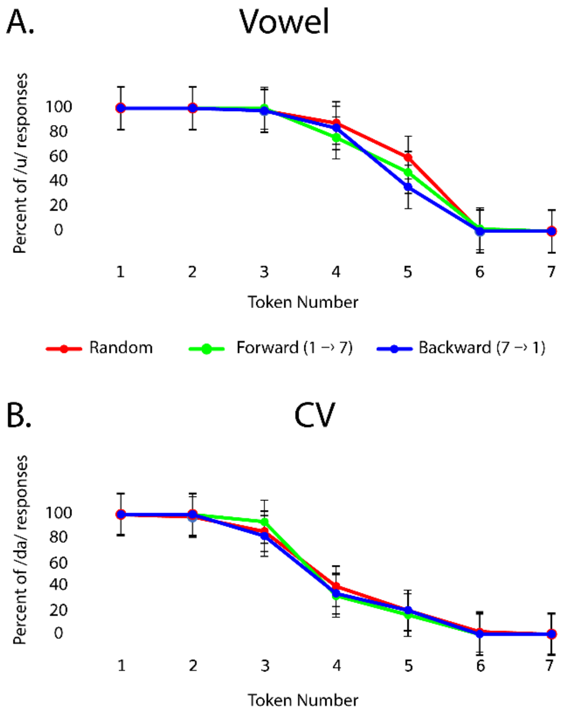


**Figure S1:** Psychometric functions (n=5), comparing the identification for **(A)** vowels and **(B)** consonant-vowel syllables (CVs). Vowels exhibited more nonlinear response patterns than CVs as evidenced by the more salient movement of the perceptual boundary (e.g., see Tk4-Tk5). Error bars = ±1 s.e.m.

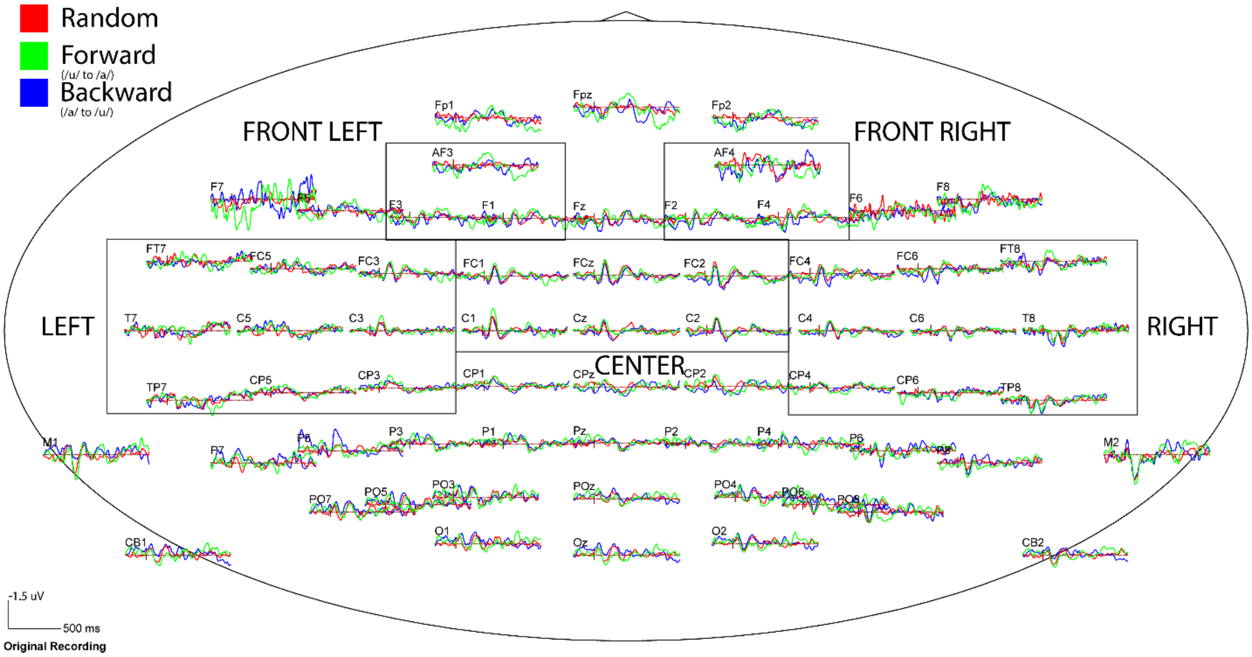


**Figure S2:** Scalp topography of ERPs. Boxes denote superchannels used in the primary analysis. Negative is plotted up.

| Subject Num | Tk3 Yule's Q | Tk5 Yule's Q | Response Pattern (listener type) |
|---|---|---|---|
| S1[†] | -0.87* | -0.75* | Hysteresis |
| S2[†] | 0.08 | 0.66* | Enhanced Contrast |
| S3 | 0.00 | 0.38* | Enhanced Contrast |
| S4[†] | 0.00 | 0.00 | Critical Boundary |
| S5 | -0.84* | -0.49* | Hysteresis |
| S6 | -0.77* | 0.00 | Hysteresis |
| S7 | -1.00* | 0.00 | Hysteresis |
| S8 | -1.00* | -0.74* | Hysteresis |
| S9 | 0.00 | -0.59* | Nil |
| S10 | -0.44* | 0.00 | Hysteresis |
| S11 | -0.76* | -0.48* | Hysteresis |
| S12 | -0.23 | 0.40* | Enhanced Contrast |

| | | | |
|---|---|---|---|
| S13 | -0.60* | -0.30 | Hysteresis |
| S14 | 0.00 | 0.88* | Enhanced Contrast |
| S15 | -0.76* | -0.17 | Hysteresis |

**Table S1:** Yule's Q values for Tk3/5 (i.e., tokens flanking the expected β0) and response patterns by participant. More negative/positive Yule's Q denotes hysteresis/enhanced contrast response patterns, respectively. *Yule's Q of medium-to-large effect size |Q| ≥ 0.33. †Individuals shown in **Fig. 1E-G**
